# Validation and Comparison of a Modified CDC Assay with two Commercially Available Assays for the Detection of SARS-CoV-2 in Respiratory Specimen

**DOI:** 10.1101/2020.06.29.179192

**Authors:** Amorce Lima, Vicki Healer, Elaine Vendrone, Suzane Silbert

**Author notes:** Corresponding author Amorce Lima, PhD, Esoteric Testing Laboratory, Tampa General Hospital, 1 Tampa General Circle, Tampa, FL. USA. Zip code: 33606-3571.

## Abstract

Severe acute respiratory syndrome coronavirus 2 (SARS-CoV-2), the virus that causes coronavirus disease 2019 (COVID-19), has spread rapidly around the globe since it was first identified in December of 2019 in Wuhan, China. In a race to contain the infection, researchers and healthcare officials have developed several assays to help diagnose individuals with COVID-19. To help laboratories in deciding what assay to bring into testing lines, factors such as assay availability, cost, throughput, and TAT should be considered. Here we validated a modified version of the CDC assay and used it as a reference to evaluate the performance of the NeuMoDx™ SARS-CoV-2 and DiaSorin Simplexa™ Covid-19 Direct assays. *In silico* analysis and clinical sample testing showed that the primesr/probes designed by the CDC were specific to the SARS-CoV-2 as they accurately detected all reactive samples with an assay LoD of 200 copies/ml. The performance of the three assays were analyzed using 161 nasopharyngeal swabs specimen tested within 24 hours or 5 days from routine testing. A 100% agreement was observed between the commercial assays and the modified CDC SARS-CoV-2 assay. A deeper look at the Ct values showed no significant difference between NeuMoDx and the modified CDC SARS-CoV-2 assay, whereas DiaSorin had lower overall Ct values than the modified CDC SARS-CoV-2 assay. NeuMoDx and DiaSorin workflows were much easier to perform. NeuMoDx has the highest throughput and shortest TAT, whereas although the modified CDC SARS-CoV-2 assay has comparable throughput to DiaSorin, it has the longest hands-on time, and highest TAT.

## INTRODUCTION

Coronaviruses are a family of enveloped positive-RNA viruses that infect vertebrates and can widely spread among human causing diseases in the respiratory, intestinal, liver and nervous systems (1). Some coronaviruses, namely human coronaviruses OC43, 229E, NL63 and HKU1, have been recognized as the etiological agents of common cold. However, others have been associated with severe respiratory illness and outbreaks: Severe acute respiratory syndrome coronavirus (SARS-CoV) resulted in global outbreak in 2002, the Middle East Respiratory Syndrome (MERS-CoV) appeared in 2012, and SARS-CoV-2 of 2019 pandemic (2-5). SARS and MERS are closely-related coronaviruses of about 30 kb genome and members of the beta coronavirus clade (5); like MERS-CoV, SARS-CoV-2 is believed to have been originated from bats. Symptoms associated with COVID-19, the disease caused by SARS-CoV-2, vary from mild to severe illness depending on the individual’s underlying medical conditions. They include cough, shortness of breath, fever, chills, and muscle pain (https://www.cdc.gov/coronavirus/2019-ncov/symptoms-testing/symptoms.html).

SARS-CoV-2, which started in Wuhan City in China at the end of December 2019, has spread to over 188 countries and was declared a global pandemic by the World Health Organization (WHO) on March 11^th^ (6). As of June 26, 2020, there were more than 9,680,000 confirmed cases of COVID-19, and 491,000 deaths due to the outbreak (https://coronavirus.jhu.edu/map.html). The United States lead the world with over 2,446,000 cases of COVID-19 and over 125,000 deaths as of June 26, 2020. The rate of new cases is increasing in the United States, and in countries such as Mexico and Brazil with no apparent end in sight. In addition to the health crisis caused by this pandemic, the economic impact is equally devastated with the world facing the worst recession since the Great Depression of the 1930s. Currently, except for Remdesivir which has been authorized by the Food and Drug Administration (FDA) for emergency use in the USA for Covid-19 treatment, there is no other therapeutics or vaccine approved from treatment of COVID-19.

There has been a race against time to develop tests for SARS-CoV-2 detection so individuals with COVID-19 could be identified and isolated to slow the spread of the disease. In January 2020, the CDC developed a TaqMan probe based molecular test, and at the end of February of the same year, the FDA provided guidance for validation of laboratory develop tests (LDTs) and submission for Emergency Used Authorization of such tests (7). In the following month, several commercially available assays became available and have been used in the laboratory under the FDA’s EUA (8-12). With the rapid development of those tests, came the challenge of assay sensitivity and specificity. The molecular assays for detection of SARS-CoV-2 are based on reverse transcriptase polymerization chain reaction (RT-PCR) targeting the open reading frame (ORF1ab), RNA dependent RNA polymerase (RdRp) and genes that encode structural proteins such as nuclear capsid (N), spike (S), envelop (E), and membrane (M) (7, 13, 14). It has been reported that the CDC N2 and WHO E-gene primer/probe sets are among the most sensitive primer/probe sets for SARS-CoV-2 detection (15).

Our laboratory at Tampa General Hospital validated a modified version of the CDC assay following the FDA EUA guidelines and brought in commercial assays under the FDA EUA to help respond to the testing demand in our hospital. All three assays are based on real-time RT-PCR to detect the viral RNA. The modified CDC SARS-CoV-2 assay involves an off-instrument cell lysis step, a nucleic acid extraction step and an amplification step on a different instrument. The assay includes a panel of two primer/probe sets targeting the viral N gene. The NeuMoDx™ SARS-CoV-2 assay and DiaSorin Simplexa™ Covid-19 Direct assays are automated sample-to-answer assays. They are both multiplex assays targeting two different regions of the viral genome. NeuMoDx targets the Nsp2 and the N genes while DiaSorin targets the ORF1ab and S gene. In this study, we sought to describe the modified CDC SARS-CoV-2 assay and compare its performance and workflow to that of the NeuMoDx™ SARS-CoV-2 and DiaSorin Simplexa™ Covid-19 Direct assays.

## MATERIALS AND METHODS

### Primers and probes

The primer/probe sets which were designed and described in the CDC protocol for the detection of SARS-CoV-2 were used in this study (Table 1) (16). N1 and N2 amplified regions of the nucleocapsid gene of the virus. An additional primer/probe set to detect the human ribonuclease (RNase) P gene (RP) in clinical specimens was also included in the panel. An *in silico* analysis of the primer and probe sequences was conducted based on the currently available sequences in the National Center for Biotechnology Information (NCBI) database as of March 11, 2020. SARS-CoV-2 RNA (strain USA_WA1/2020), 2019-nCoV_N_Positive Control and Hs_RPP30_Positive Control plasmids were included in the study as templates and external controls. SARS-CoV-2 RNA was kindly provided by the University of Texas Medical Branch (UTMB) in Galveston while the plasmids and primer/probe sets were acquired from Integrated DNA Technologies (IDT) (Coralville, USA).

**Table 1.**
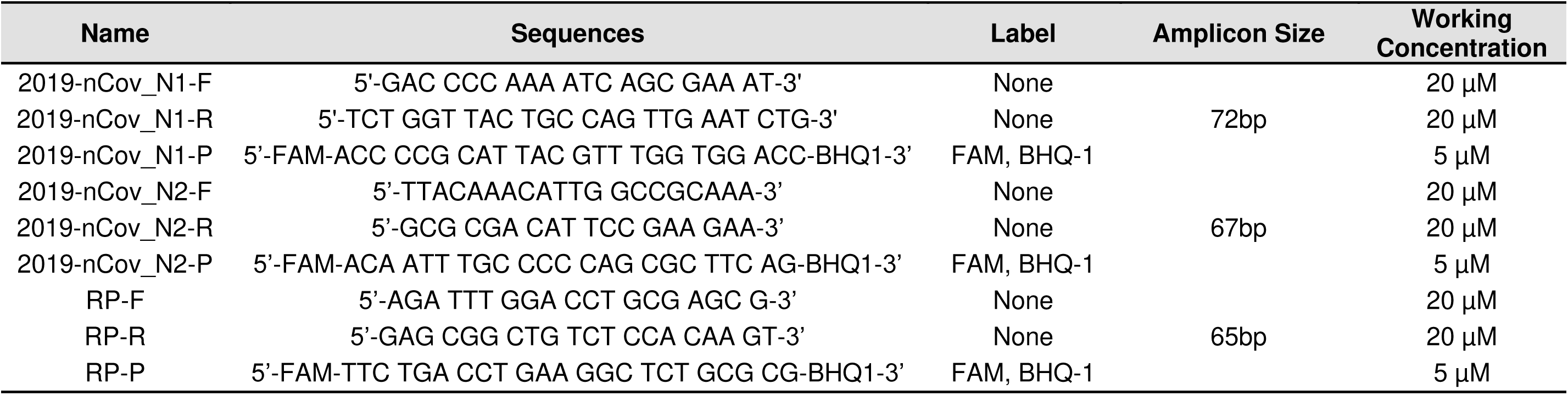
CDC primer/probe panel for RT-PCR detection of SARS-CoV-2 and human RNase P gene internal control

### Specimen enrollment

Residual respiratory specimens submitted for Covid-19 testing at Tampa General Hospital (TGH) Esoteric Testing Laboratory were used in this study according to the standard procedures and in compliance with the TGH Institutional Review Board (IRB) protocols. Those specimens included sputum, nasopharyngeal (NP) or nasal swabs in Copan Universal Transport Media (UTM) (Copan Diagnostics, Murrieta, CA) or BD Universal Viral Transport (UVT) system (BD Diagnostics, Sparks, MD). Following routine patient testing, specimens were kept at 2-8°C and tested within 24 hours or 5 days.

### Sample processing

For the modified CDC SARS-CoV-2 assay, total nucleic acid was extracted from 200 μl of sample on the bioMérieux NucliSens® easyMAG® automated system (bioMerieux, France) and eluted in 50 μl of EasyMag elution buffer. A separate reaction mix containing 5 μl of the eluate, 5 μl of TaqPath™ 1-Step RT-qPCR Master Mix (4x), 1.5 μl of combined primer/probe Mix (500nM and 125nM final concentration of primers and probes, respectively), 13.5 μl of Nuclease-free Water was made for each of the assay target (N1, N2, and RP). A no-template water control (NTC) and 2019-nCoV_N_ Positive Control (nCoVPC) were used as template for each of the primer/probe set; Hs_RPP30_Positive Control plasmid was used as template for RP primer/probe where necessary. The real-time reverse transcriptase PCR (rRT-PCR) cycling conditions were set up on the Rotor-Gene 3000 thermocycler (Corbett Research, Australia) as followed: 25°C for 2 min, 50°C for 15 min, 95°C for 2 min, followed by 45 cycles of 95°C for 3 s and 55°C for 30 s with fluorescence (FAM) detection during the 55°C incubation step. A sample was positive for SARS-CoV-2 if at least one of the two targets (N1, N2) was detected regardless whether RP was amplified, negative if none of the targets was detected and RP was detected, and invalid if RP and the two targets were not detected.

Samples were processed for the NeuMoDx™ SARS-CoV-2 assay and the DiaSorin Simplexa™ Covid-19 Direct assay according the manufacturer’s procedures. For NeuMoDx™ SARS-CoV-2 assay, 400 μl of sample was mixed with 400 μl of NeuMoDx Viral Lysis Buffer in a secondary tube before loading it onto the NeuMoDx(tm) 96 Molecular System (NeuMoDx, Ann Arbor, MI). The NeuMoDx(tm) 96 Molecular System is a sample in results out random-access platform that incorporates extraction, real-time PCR, and signal detection and analysis. A sample was positive if either or both N or Nsp2 genes were detected, negative if both targets were not amplified and the sample processing control (SPC2) was amplified, and indeterminate or unresolved if there was an instrument error or it failed to produce a valid result. For the DiaSorin Simplexa™ Covid-19 Direct assay, 50 μl of sample and 50 μl of the Reaction Mix were loaded to each well and processed according to the manufacturer’s protocol on the LIASON® MDX instrument (DiaSorin, Saluggia, Italy). Upon completion of the run, results were automatically analyzed by the LIAISON® MDX Studio software. A sample was positive for SARS-CoV-2 if either ORF1ab or S gene was detected, negative if both targets were not amplified and the internal control was amplified, and invalid if there was an internal control failure.

### Analytical sensitivity and specificity of the modified CDC SARS-CoV-2 assay

A series of two-fold dilutions of SARS-CoV-2 strain USA_WA1/2020 RNA were spiked in pooled sputum at concentrations of 800 copies/ml to 0.05 copy/ml in order to determine the limit of detection (LoD) of the assay. All samples were processed and tested in triplicate as described above. The LoD was confirmed by further testing in 20 replicates. The analytical specificity was determined by testing 22 samples which include 15 patient samples, 4 ATCC strains, and 3 commercially available nucleic acid controls.

### Clinical evaluation of the modified CDC SARS-CoV-2 assay

The performance of the modified CDC SARS-CoV-2 assay was established by testing 30 contrived NP swabs and sputum specimens and 30 non-reactive specimens. Of the 30 contrived specimens, 20 were spiked with SARS-CoV-2 strain USA_WA1/2020 RNA at 1x-2x LoD concentrations and 10 were spiked at concentrations spanning the assay’s testing range (60,000 copies/ml to 234 copies/ml).

### Performance comparison between NeuMoDx SARS-CoV2 assay, the Simplexa Covid-19 Direct assay and the modified CDC SARS-CoV-2 assay

A total of 161 NP swabs were used to compare the clinical performance of two commercially available assays against the modified CDC SARS-CoV-2 assay. Of those, 118 samples were tested to compare NeuMoDx SARS-CoV-2 assay with the modified CDC SARS-CoV-2 assay, and 43 were tested to compare Simplexa Covid-19 Direct assay with the modified CDC SARS-CoV-2 assay.

### Statistical analysis

The results obtained from each assay were compared with those obtained using the modified CDC SARS-CoV-2 assay. EP Evaluator was used to calculate positive percent agreement (PPA), negative percent agreement (NPA), and Cohen’s kappa (k) with 95% confidence intervals.

## RESULTS

### Primer/probe analysis and PCR efficiency testing

The primer/probe sets used in the modified CDC SARS-CoV-2 assay were designed by the CDC (16). Since the CDC primer/probe panel became available, many SARS-CoV-2 isolates have been sequenced. Therefore, we conducted an analysis based on the currently available sequences in the National Center for Biotechnology Information (NCBI) database as of March 11, 2020. Primer Blast analysis showed that the primer/probe sets were specific to all available sequences for SARS-CoV-2. Multiple sequence alignment (CLUSTAL multiple sequence alignment by MUSCLE) of the nucleocapsid gene of SARS-CoV-2 and the two closely related coronaviruses, SARS-CoV and MERS-CoV, showed the positions of the primer/probe sets (Fig. 1A). The N1 set amplified a 72 bp fragment of the nucleocapsid gene, N2 amplified a 67 bp of the same gene, and the primer/probe set for human RP gene internal control produced a 57 bp amplicon (Fig. 1B).

**Figure 1.**
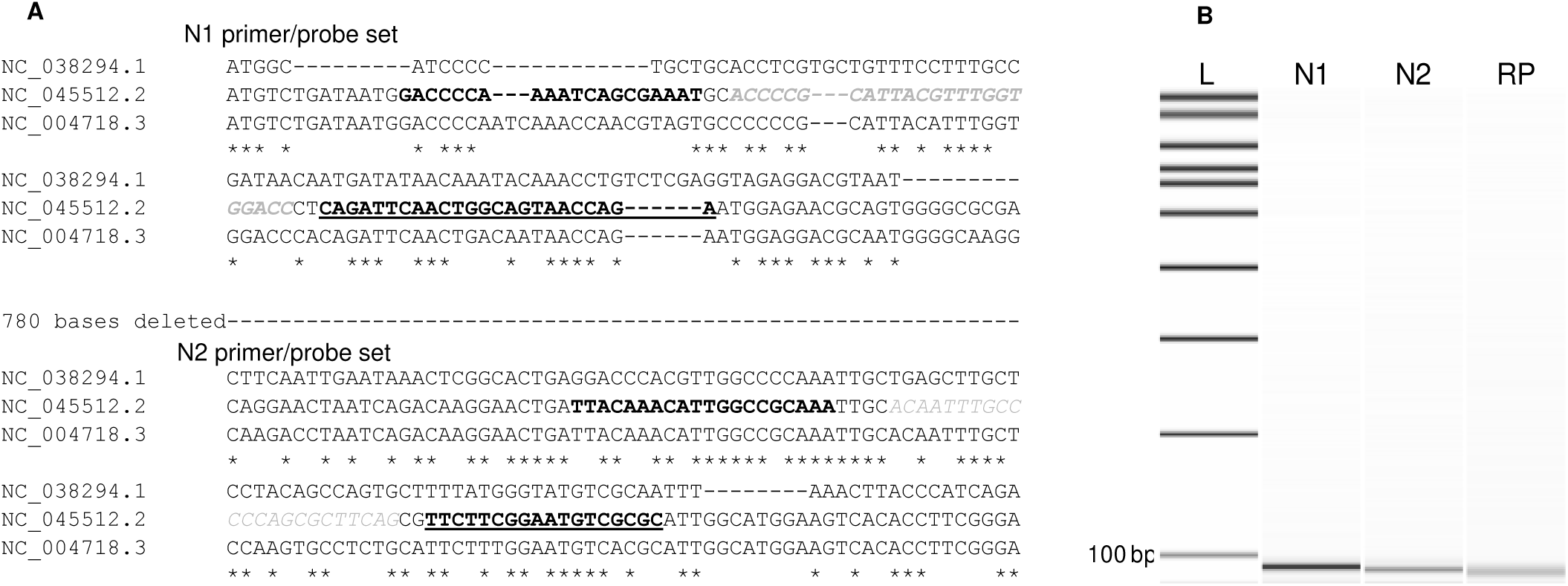
A) Multiple sequence alignment of partial sequence of the nucleocapsid gene of MERS-CoV (NC 038294.1), SARS-CoV-2 (NC_045512.2), SARS-CoV (NC_004718.3) showing target regions of N1 and N2 primer/probe sets. Forward primers sequences are bolded, probes are faded and itilicized, and reverse primers are underlined and bolded. Note: 780 bases between the two primer sets were omitted to shorted the length of the sequence. B) Capilary gel electrophoresis picture generated using the Agilent DNA 7500 kit on the Agilent 2100 bioanalyzer instrument. Image showed single band for each primer set. From left to right: L (ladder:100 – 7000bp), N1, N2, and RP of approximately 72 bp, 67 bp, and 65 bp, respectively.

The PCR efficiency was determined for each of the primer/probe set by testing a series of 10-fold dilutions of the 200,000 copies/μl concentration of the nCoVPC plasmid. The data showed that the PCR was linearity over 6 orders of magnitude with great PCR efficiency (N1=111% and N2=100) and R^2^ of 0.99 for each of the primer/probe set (Fig. 2). The standard curve generated using the known concentrations was used to accurately determine the concentration of the USA_WA1/2020 RNA. It was found that the USA_WA1/2020 RNA obtained from UTMB was at a concentration of 1.5 × 10^8^ copies/ml.

**Figure 2.**
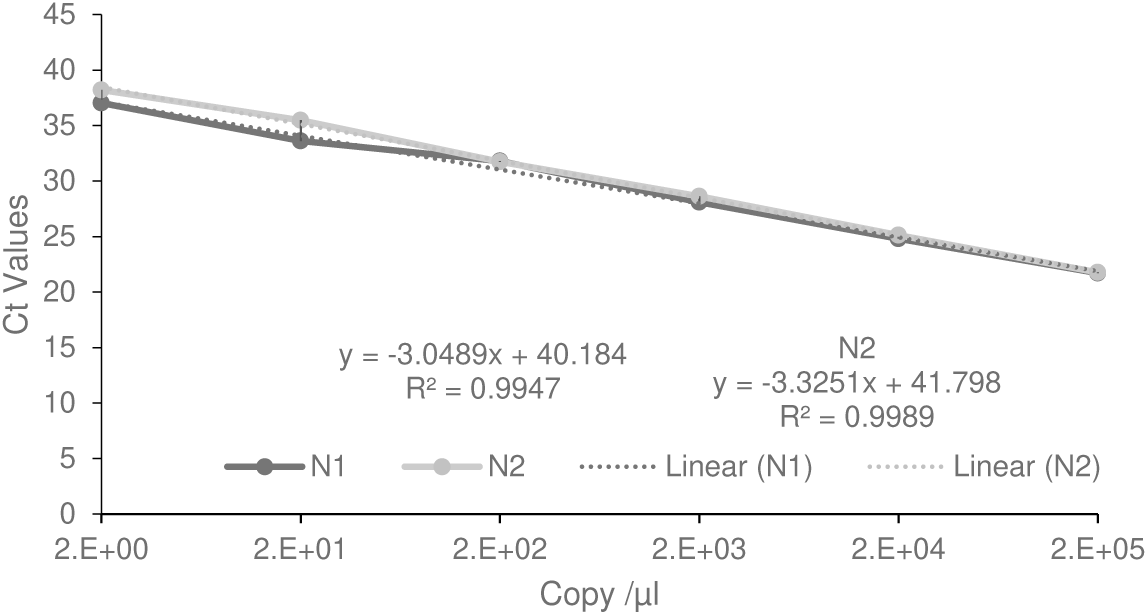
Real time PCR determining amplification efficiency of the primer/probe sets. Ten-fold serial dilution of 200,000 copies/μl of nCoVPC plasmid was tested. PCR linearity over 6 orders of magnitude with a limit of detection of 2 copies/μl; N1 slope of −3.05 with a correlation coefficient R2 = 0.99; N2 slope =-3.33 and R2 = 0.99.

### Analytical sensitivity and specificity of the modified CDC SARS-CoV-2 assay

The analytical sensitivity of the modified CDC SARS-CoV-2 assay was determined by testing pooled sputum samples spiked with a series of two-fold dilutions at concentrations of 800 copies/ml to 50 copy/ml of SARS-CoV-2 strain USA_WA1/2020. The lowest concentration that produced a threshold cycle (Ct) value for all three replicates was determined to be the LoD; it was estimated to be 200 copies/ml (Table 2A). The LoD was confirmed by further testing 20 replicates that were inoculated with 200 copies/ml; all 20 replicates were tested positive (Table 2B). The analytical specificity of the assay was determined by testing 22 samples, which included 15 patient samples, 4 ATCC strains, and 3 commercially available nucleic acid controls. The result showed that none of the targets was detected (Table 3).

**Table 2.**
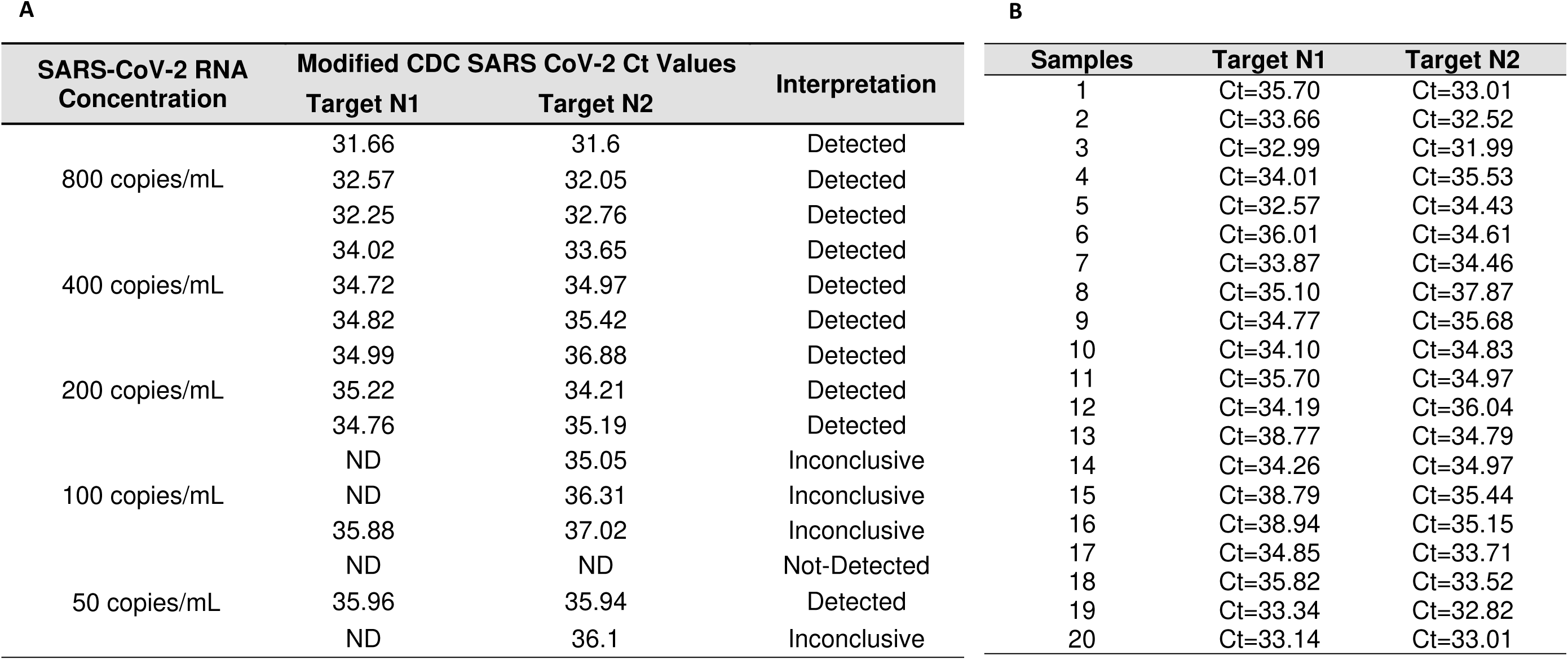
Limit of detection of the modified CDC SARS-CoV-2 assay

**Table 3.** Proficiency panel of viral and bacterial samples tested using the modified CDC SARS-CoV-2 assay

| Organisms | Source | Method of Identification | Interpretation |
| --- | --- | --- | --- |
| MERS CoV | Control Suspension | IDT# 10006623 | Not-Detected |
| SARS CoV | Control Suspension | IDT# 10006624 | Not Detected |
| Adenovirus | Control Suspension | Exact DX# ADVH102 | Not-Detected |
| Coronavirus HKU1 | NP Patient Sample | BioFire RVP2 | Not-Detected |
| Coronavirus NL63 | NP Patient Sample | BioFire RVP2 | Not-Detected |
| Coronavirus 229E | NP Patient Sample | BioFire RVP2 | Not-Detected |
| Coronavirus OC43 | NP Patient Sample | BioFire RVP2 | Not-Detected |
| Influenza A | NP Patient Sample | BioFire RVP2 | Not-Detected |
| Influenza B | NP Patient Sample | BioFire RVP2 | Not-Detected |
| Respiratory Syncytial Virus | NP Patient Sample | BioFire RVP2 | Not-Detected |
| Human Metapneumovirus | NP Patient Sample | BioFire RVP2 | Not-Detected |
| Human Rhinovirus | NP Patient Sample | BioFire RVP2 | Not-Detected |
| *Chlamydia pneumoniae* | NP Patient Sample | BioFire RVP2 | Not-Detected |
| *Mycoplasma pneumoniae* | NP Patient Sample | BioFire RVP2 | Not-Detected |
| Parainfluenza Virus 1 | NP Patient Sample | BioFire RVP2 | Not-Detected |
| Parainfluenza Virus 2 | NP Patient Sample | BioFire RVP2 | Not-Detected |
| Parainfluenza Virus 3 | NP Patient Sample | BioFire RVP2 | Not-Detected |
| Parainfluenza Virus 4 | NP Patient Sample | BioFire RVP2 | Not-Detected |
| *Staphylococcus epidermidis* | 0.5 McFarland Suspension | ATCC Strain 14990 | Not-Detected |
| *Streptococcus pyogenes* | 0.5 McFarland Suspension | ATCC Strain 19615 | Not-Detected |
| *Pseudomonas aeruginosa* | 0.5 McFarland Suspension | ATCC Strain 27853 | Not-Detected |
| *Mycobacterium tuberculosis* | 0.5 McFarland Suspension | ATCC Strain 25177 | Not-Detected |

### Clinical evaluation of the modified CDC SARS-CoV-2 assay

Since there was no available positive COVID-19 specimen at the time, samples were inoculated with a wide range of SARS-CoV-2 strain USA_WA1/2020 RNA to mimic the extent of viral colonization in respiratory specimen. The performance of the assay was evaluated in 60 sputum samples (30 contrived positives and 30 negatives). Of the 30 contrived positive samples, 19 were spiked with 200 to 400 copies/ml of SARS-CoV-2 and one known positive patient sample; the remaining 10 were spiked with 234 copies/ml to 60,000 copies/ml. The results showed that the assay detected SARS-CoV-2 in all 30 reactive samples, whereas no amplification was seen in the 30 non-reactive samples (Table 4).

**Table 4.**
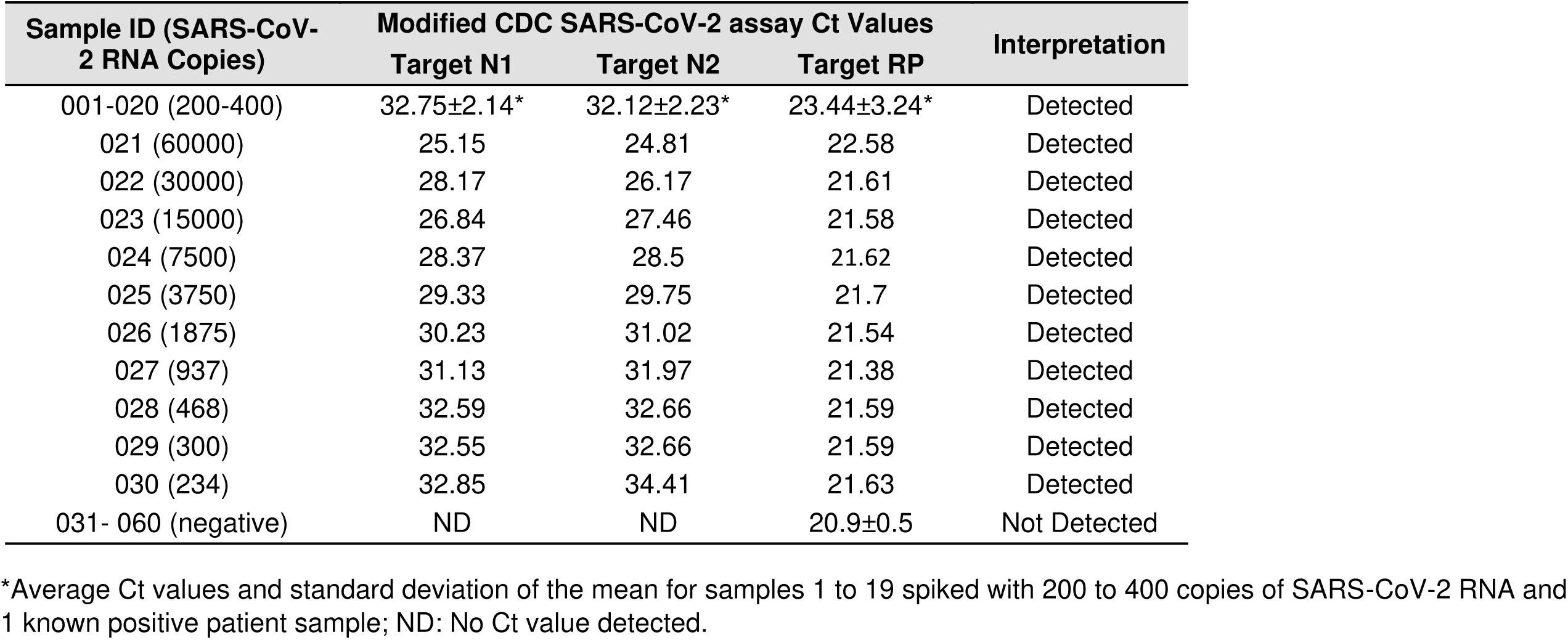
Ct values of contrived clinical specimen spiked with different concentrations of SARS-CoV-2 RNA

| Sample ID (SARS-CoV-2 RNA Copies) | Modified CDC SARS-CoV-2 assay Ct Values |  |  | Interpretation |
| --- | --- | --- | --- | --- |
|  | Target N1 | Target N2 | Target RP |  |
| 001-020 (200-400) | 32.75±2.14* | 32.12±2.23* | 23.44±3.24* | Detected |
| 021 (60000) | 25.15 | 24.81 | 22.58 | Detected |
| 022 (30000) | 28.17 | 26.17 | 21.61 | Detected |
| 023 (15000) | 26.84 | 27.46 | 21.58 | Detected |
| 024 (7500) | 28.37 | 28.5 | 21.62 | Detected |
| 025 (3750) | 29.33 | 29.75 | 21.7 | Detected |
| 026 (1875) | 30.23 | 31.02 | 21.54 | Detected |
| 027 (937) | 31.13 | 31.97 | 21.38 | Detected |
| 028 (468) | 32.59 | 32.66 | 21.59 | Detected |
| 029 (300) | 32.55 | 32.66 | 21.59 | Detected |
| 030 (234) | 32.85 | 34.41 | 21.63 | Detected |
| 031- 060 (negative) | ND | ND | 20.9±0.5 | Not Detected |

\*Average Ct values and standard deviation of the mean for samples 1 to 19 spiked with 200 to 400 copies of SARS-CoV-2 RNA and 1 known positive patient sample; ND: No Ct value detected.

### Clinical performance comparison of the NeuMoDx SARS-CoV-2 assay, Simplexa Covid 19 Direct assay compared to the modified CDC SARS-CoV-2 assay

One hundred and sixty-one NP samples including those with a wide range of Ct values were used to compare the performance of the three assays. Of those, 67 positive and 51 negative samples were used to compare NeuMoDx SARS-CoV-2 assay with the modified CDC SARS-CoV-2 assay, while 22 positive and 21 negative samples were used to compare DiaSorin Simplexa Covid 19 Direct assay with the modified CDC SARS-CoV-2 assay. For comparison between modified CDC SARS-CoV-2 assay and Simplexa Covid 19 Direct assay, 37 samples were run within 2 days and 6 were run within 5 days of first testing. For comparison between the modified CDC SARS-CoV-2 assay and NeuMoDx SARS-CoV-2 assay, 104 samples were run within a day and 14 were run within 5 days of first testing. All 89 positive and 72 negative samples tested by the modified CDC SARS-CoV-2 assay matched the results using the two commercial assays evaluated yielding a 100% PPV and NPV for each assay (Table 5A). Two samples that were negative on the modified CDC SARS-CoV-2 assay resulted in indeterminate on the NeuMoDx for the Nsp2. NeuMoDx did not yield a Ct value for one of the two targets in 5 samples: 4 were negative for Nsp2 gene target and 1 was negative for the N gene target. However, they still considered positive as only one detected target was needed for a positive result.

**Table 5.** Clinical performance and workflow comparison between modified CDC SARS-CoV-2 assay and the commercial assays

| A |  |  |  |  |  |  | B |  |  |  |  |
| --- | --- | --- | --- | --- | --- | --- | --- | --- | --- | --- | --- |
| Commercial Assays |  | Modified CDC SARS-CoV-2 Assay |  |  | Kappa (%) | PPA/NPA (%) | 95% CI | *Workflow | Assays |  |  |
|  |  | Positive | Negative | Total |  |  |  |  | Modified CDC Assay | NeuMoDx | DiaSorin |
| NeuMoDx | Positive | 67 | 0 | 69 | 100 | 100/100 | (96.9-100) | Hands-on | ~45min | ~5min | ~15min |
|  | Negative | 0 | 51 | 51 |  |  |  | Extraction | ~35min | ~1h20min | N/A |
|  | Total | 67 | 51 | 118 |  |  |  | PCR | ~90min |  | ~1h25min |
| DiaSorin | Positive | 22 | 0 | 22 | 100 | 100/100 | (91.8-100) | Overall TAT | ~2h50min | ~1h25min | ~1h40min |
|  | Negative | 0 | 21 | 21 |  |  |  | Max samples/run | 10 | 24 | 8 |
|  | Total | 22 | 21 | 43 |  |  |  | **Throughput/8-h shift | ~40 | ~96 | ~40 |

PPA: Positive percent agreement; NPA negative percent agreement; CI: confidence interval
\*Workflow and overall turnaround time (TAT) is based on 8 samples per run; \*\* Number of samples that can be resulted in an 8-h shift.

Although 100% agreement was observed among the assays, further analysis showed that there were some differences in the Ct values for each assay. The lowest and highest Ct values recorded for the modified CDC SARS-CoV-2 assay were 12.24 and 38.52; for the NeuMoDx SARS-CoV-2 assay 11.45 and 36.65, and for DiaSorin Simplexa Covid 19 Direct assay 12.0 and 34.3. The average Ct value difference in samples run within 24 hours between NeuMoDx SARS-CoV-2 and the modified CDC SARS-CoV-2 assay was −0.14, and −2.13 between samples run within 5 days. The overall Ct value difference for all samples run between the two assays was −0.346. On the other hand, the average Ct values difference between samples run within 2 days between DiaSorin Simplexa Covid 19 Direct assay and the modified CDC SARS-CoV-2 assay was −2.42, and −6.0 between samples run within 5 days. The overall Ct value difference for all samples between the two assays was −3.47.

Apart from the clinical performance evaluation, we also assessed the workflow of each assay and compared the time of sample-to-result for each platform. Based on 8 samples/run, it was estimated to be 2 h and 50 min for modified CDC SARS-CoV-2 assay, which included a 35 min extraction time, 90 min PCR run, and 45 min of hands-on time. It was estimated to be 1 h and 40 min including 15 min hands-on time for DiaSorin Simplexa Covid 19 Direct assay. The NeuMoDx SARS-CoV-2 assay resulted samples in 1 h and 25 min which included a 5 min hands-on time (Table 5B). The above estimation was based on 8 samples due to number of samples that could be loaded on the DiaSorin instrument per run. However, the NeuMoDx instrument is a random-access platform that can process and result 24 samples at a time in less than two hours. That enabled the NeuMoDx assay to have a throughput of 96 samples per 8-hr shift compared to 40 samples on the DiaSorin Simplexa Covid 19 Direct assay, and 40 samples on the modified CDC SARS-CoV-2 assay if sample processing and PCR runs were staggered.

## DISCUSSION

It has been three months since the WHO declared SARS-CoV-2 a global pandemic. Although the overall infection and death rates of COVID-19 has been declining in some countries, it has increased in others. Therefore, the ongoing pandemic still poses great risks for many around the world, and with the easing of certain restrictions, the need for health care facilities to be equipped and accurately test for the virus to limit its spread is as crucial as it will ever be. To that extent, laboratories have brought in SARS-CoV-2 assays and molecular platforms to respond to the need of their communities. There have been a few publications on head-to-head comparisons of those assays, including a couple very recently as we preparing this article, in order to shed light on their performance characteristics and help laboratories make informed decisions on acquiring those assays (9, 17-19).

In this study, we validated a modified CDC SARS-CoV-2 assay and compared its performance to two commercial automated sample-to-answer assays for the detection of SARS-CoV-2 RNA. We confirmed that the primer/probe sets were specific to all SARS-CoV-2 clades based on the available genome sequences including those that were not available at the time when those primer sets were originally designed. Not only were those primers/probe sets specific to SARS-CoV-2 based on *in silico* study, there were no false positive results in cross-reactivity experiments using a panel of bacterial and closely-related virus targets. We found that the primer/probe sets have high PCR efficiency and a LoD of 200 copies/ml. The clinical sensitivity and specificity of the assay was also evaluated in samples with different concentrations of viral RNA. The results showed that the assay is specific to SARS-CoV-2 as it was only detected in the reactive samples.

The other objective of this study was to compare the clinical performance of two commercially available assays in out laboratory, NeuMoDx SARS-CoV-2 assay and Simplexa Covid-19 Direct assay, to the modified CDC SARS-CoV-2 assay. There was an overall agreement of 100% between the results obtained on the commercial assays and those on the modified CDC SARS-CoV-2 assay. It is worth noting that while we found that the modified CDC SARS-CoV-2 assay has an LoD of 200 copies/ml whereas the reported LoD in the package inserts For NeuMoDx SARS-CoV-2 assay and DiaSorin Simplexa Covid 19 Direct is 250 copies/ml and 500 copies/ml for assay, respectively.

A closer look at the Ct value differences between the modified CDC SARS-CoV-2 assay and the commercial assays suggests that there is not a significant difference between the modified CDC SARS-CoV-2 assay and NeuMoDx SARS-CoV-2; however, there seems to be a greater difference in Ct values between DiaSorin Simplexa CoV-2 Direct assay and the modified CDC SARS-CoV-2 assay, with DiaSorin having lower Ct values. The difference is even greater in samples that were run 5 days after the routine testing on the modified CDC SARS-CoV-2 assay. This is in line with previously published data that showed Ct values on DiaSorin was much lower than those on an the modified CDC SARS-CoV-2 assay by an average Ct difference of −2.1 (18). The overall data also suggest that depending on the viral burden in the samples NP samples can be refrigerated for at least 5 days and still maintain the RNA integrity for viral detection by the assays in this study.

A limitation of this study was that the same samples were not tested by the three different assays, so a head-to-head comparison of the three assays was not performed. This was due to the limit of available kits for routine testing in patient care. However, we did a head-to-head comparison of the assay’s workflow. The NeuMoDx 96 molecular system is a sample-to-answer and random-access platform that automatically performs nucleic acid extraction, amplification, and signal detection and analysis requiring only very little human interaction for loading and scanning the samples (20). It has the shortest TAT and highest throughput of the three assays. DiaSorin Simplexa CoV-2 Direct assay involves a simple operation procedure that does not include an extraction step; it has, however, much lower throughput than NeuMoDx. The modified CDC SARS-CoV-2 assay is singleplex; each target must be run in different tubes, as opposed to the two commercial assays, which are multiplex. It requires separate extraction steps on the EasyMag which can extract up to 24 samples at a time, longer hands-on time, and has much lower throughput compared to NeuMoDx. It has comparable throughput to DiaSorin if nucleic acid extraction and PCR reaction tubes for the next sample batch are prepared before the PCR cycles of the previous batch ends. Therefore, although the three assays have the same accuracy, the overall workflow favors the commercial platforms.

In conclusion, diagnostic laboratories around the world have faced with unprecedented challenges due to the SARS-CoV-2 pandemic. Thousands of SARS-CoV-2 tests are being executed in any given laboratory each day. This testing requirement has not only forced laboratory to bring in new technologists to help with testing, but it has also led to the shortage of testing reagents. Consequently, laboratories had to acquire different assays and platforms to meet testing demand. As much as LDT assays, such as the modified CDC SARS-CoV-2 assay, were instrumental at the onset of the pandemic for COVID-19 testing, their overall testing capacity are limited. Therefore, it is necessary for laboratories to acquire multiple high throughput automated instruments that can test high number of samples quickly with almost the same number of qualified laboratory professionals.

## ACKNOWLEDGEMENTS

We are grateful to University of Texas Medical Branch for providing SARS CoV-2 RNA Template for the validation of the modified CDC SARS-CoV-2 assay.

